# SUV39H1 mediated regulation of KLF4 and KDM4A coordinate smooth muscle cell phenotypic plasticity

**DOI:** 10.1101/2024.11.03.621725

**Authors:** Payel Chatterjee, Raja Chakraborty, Ashley Sizer, Brendan J. O’Brien, Yi Xie, John Hwa, Kathleen A. Martin

## Abstract

**Background:** Reversible DNA methylation contributes to the phenotypic plasticity of vascular smooth muscle cells (VSMCs). This plasticity contributes to vascular growth remodeling, but also underlies cardiovascular pathologies, including intimal hyperplasia. We investigated the role of SUV39H1, a histone methyltransferase that generates the H3K9me3 repressive epigenetic mark, in VSMC plasticity

**Methods:** We applied knockdown, qPCR, western blotting, chromatin immunoprecipitation (ChIP) assays, and RNA-Seq in human coronary artery SMCs (hCASMCs), and murine carotid ligation to determine the role of SUV39H1 in VSMC plasticity.

**Results:** Expression of SUV39H1 and the H3K9me3 mark it generates increase, whereas the cognate H3K9me3 demethylase KDM4A decreases, over time during the progression of murine intimal hyperplasia following carotid artery ligation, with marked elevation of SUV39H1 and H3K9me in the neointima. SUV39H1 knockdown induced contractile genes and contractility while decreasing migration and proliferation in hCASMCs. Transcriptomic analysis confirmed that SUV39H1 promotes SMC dedifferentiation. SUV39H1 knockdown revealed that SUV39H1 promotes KLF4 upregulation by increasing KLF4 mRNA stability. PDGF-BB induced SUV39H1 expression and SUV39H1-dependent H3K9me3 modification of contractile gene promoters in hCASMC. SUV39H1 knockdown reduced the repressive H3K9me3 and 5mC marks but increased the activating H3K27Ac mark at these promoters. SUV39H1 knockdown also increased expression of KDM4A and its binding to contractile promoters, suggesting an opposing regulatory relationship between the writer and eraser of H3K9me3.

**Conclusions:** We identify SUV39H1 as an epigenetic regulator that promotes VSMC dedifferentiation by stabilizing KLF4 expression and by altering chromatin state. We report SUV39H1-dependent dynamic regulation of the repressive H3K9me3 mark at contractile gene promoters, and opposing regulation of the enzymes that write (SUV39H1) and erase (KDM4A) these marks during VSMC phenotypic switching. These studies suggest that coordinate regulation of both histone and DNA methylation contribute to VSMC phenotypic plasticity.

**Highlights:**

- The histone methyl transferase SUV39H1 is differentially regulated in VSMC phenotypic switching in culture and in vascular remodeling. SUV39H1 and the H3K9me3 mark it generates increase, whereas the cognate H3K9me3 eraser KDM4A decreases, during the progression of murine intimal hyperplasia.
- SUV39H1 promotes PDGF-induced VSMC de-differentiation, stabilizing KLF4 mRNA. SUV39H1 opposes VSMC contractility and promotes proliferation and migration.
- PDGF-BB induces SUV39H1-dependent H3K9me3 marks at VSMC contractile gene promoters, and SUV39H1 loss of function alters multiple marks that govern chromatin accessibility at these promoters, with opposing effects on the repressive H3K9me3 and 5mC and activating H3K27Ac marks.
- SUV39H1 generates H3K9me3 marks that are associated with silenced heterochromatin. We note dynamic regulation of this mark at VSMC contractile gene promoters, and opposing regulation of the enzymes that write (SUV39H1) and erase (KDM4A) these marks during VSMC phenotypic switching.

## Introduction

Vascular smooth muscle cells (SMC) which make up the muscular layer of blood vessel wall possess a remarkable plasticity that allows them to reversibly de-differentiate and adopt alternate fates in response to environmental cues. This ability to dramatically alter their phenotype allows SMC to contribute to vascular growth, remodeling, and repair but, inappropriate SMC phenotypic modulation also underlies many cardiovascular pathologies, including atherosclerosis, restenosis, intimal hyperplasia, transplant vasculopathy, pulmonary hypertension and aneurysm^1^. Differentiated SMC are quiescent and exhibit a classical elongated myocyte morphology and express a repertoire of SMC-specific contractile genes that allow for their contractile function^2^. In response to extracellular signals including growth factors such as platelet-derived growth factor (PDGF) released at sites of injury, mature SMCs undergo phenotypic transition from a contractile, quiescent phenotype to the “synthetic” or fibroblast-like phenotype accompanied by downregulation of the classical contractile markers of SMC, an increase in extracellular matrix synthesis (collagens, fibronectin), proliferation and migration^2^. Identifying new mechanisms that underlie SMC phenotypic switching may provide fundamental understanding of the biology of cellular fate transitions and may help identify new potential targets and therapeutic strategies for cardiovascular diseases.

SMC phenotypic modulation is a highly regulated and coordinated by transcriptional and epigenetic control. Transcription factors MYOCD and SRF are recruited to conserved cis regulatory CArG elements and are essential for SMC contractile gene expression ^3^, whereas KLF4 is a master SMC de-differentiation factor^4^. Chromatin remodeling through epigenetic modifications has been shown to influence SMC phenotype^4^. Reversible DNA methylation contributes to phenotypic switching with DNA methyltransferases (DNMTs) promoting dedifferentiation ^4^. The TET2 methylcytosine dioxygenase reverses DNA methylation and serves as a master pro-differentiation mechanism in SMC ^5^. Histone acetylation is an activating modification. The histone acetyltransferases p300 and CBP, and histone deacetylases (HDACs) also play key roles in VSMC phenotypic switching ^6^.

Histone methylation is another important epigenetic regulatory mechanism controlled by the actions of methyltransferases and demethylases. Tri-methylation at Histone H3 lysine 9 (H3K9me3) is generated by histone methyl transferase SUV39H1 (suppressor of variegation 3-9 homolog 1)^7^. This mark is associated with constitutive heterochromatin and transcriptional repression^8^ . SUV39H1 is repressed in diabetic SMCs leading to de-repression of inflammatory genes ^9, 10^. The H3K9me3 epigenetic mark is reduced in SMC atherosclerotic plaques ^11^ and viral mediated silencing of SUV39H1 in rat vessels protects against intimal hyperplasia ^12^. However, the molecular mechanisms by which SUV39H1 might attenuate neointimal hyperplasia and its role in SMC plasticity remain poorly characterized. Herein, we address regulation of H3K9 trimethylation and the roles of SUV39H1 and the H3K9me3-specific demethylase KDM4A in VSMC phenotypic modulation.

## Methods

### Animal studies

All experiments were approved by the Institutional Animal Care and Use Committees of Yale University and in adherence to the NIH Guide for the Care and Use of Laboratory Animals. For carotid ligation,12-week-old male C57BL/6 mice were anesthetized with ketamine and xylazine. The left common carotid artery was exposed and completely ligated using 6-0 silk suture immediately proximal to the carotid bifurcation. Mice were euthanized and the left injured and uninjured right carotid arteries were collected at 3,7,14-and 28-days post-surgery. Tissues were fixed in 4% paraformaldehyde, transferred to 30% sucrose solution, embedded in OCT compound (Tissue Tek), frozen and stored at -80°C for cryosectioning.

### Cell culture

Human coronary artery smooth muscle cells (hCASMCs, Lonza) were propagated in M199 media (Gibco, 11150-067) supplemented with 10% FBS, 100 U/ml each penicillin-streptomycin, 2.7 ng/ml rhEGF (Biolegend, 713008), and 2ng/ml rhFGF (Biolegend 71034). Cells from passages 5 through 7 as described earlier ^5, 6^ were used for all experiments. The cells were treated with Rapamycin (Sigma, R8781) with final concentration (50 nM), or PDGF-BB (10ng/ml) (Sigma, P3201) or vehicle (ethanol for rapamycin, PBS for PDGF-BB) for indicated time points.

### Transient transfection of siRNA

hCASMCs were transfected using Lipofectamine RNAiMAX (Life Technologies, 1377850) with siRNA targeting SUV39H1 (siSUV39H1, Dharmacon, # L-009604-00-0010), final concentration 100nM, KDM4A (siKDM4A, Dharmacon, # L-004292-00-0010), final concentration 25nM (or non-silencing siRNAs (“scrambled” siControl, Dharmacon#D-001810-10-20), final concentration used 100nMBriefly, cells were plated in six-well plates or 10-cm dishes at a confluency of 50 to 60% and transfected the next day using Lipofectamine containing siRNA according to the manufacturer’s protocol. After 7 hours, media was replaced with M199 media with 10% FBS. 24 hours later, cells were serum starved for experiments as described above.

### Protein stability assay

24 hours after seeding in six-well plates at a confluency of 50 to 60%, hCASMCs were starved in 2.5% FBS for another 24 hours. The next day, cells were treated with Rapamycin at a final concentration (50nM). Following Rapamycin treatment for 24 hours, cycloheximide (Sigma, 0.1 mg/ml) was added and cells were harvested at indicated time points. For 0-hour control, untreated cells were harvested immediately before cycloheximide treatment. Cell pellets were lysed in RIPA buffer supplemented with 1x protease inhibitor (Roche, 4693124001) and phosphatase inhibitor (Roche, 4906845001) and subjected to Western blot.

### Migration assay

Boyden chamber migration assays were performed using a 24-well Trans well chamber system (Costar 3422, Corning Inc., NY, USA) and as described earlier ^6^. Cells were seeded in the upper chamber at 2×10^4^ cells/ml in 0.1 ml serum-free M199 media. Media supplemented with 10ng/ml PDGF-BB was placed in the bottom well in a volume of 0.8 ml (used as a chemoattractant). After incubation for 8 hours at 37°C, migrated cells on the lower surface were stained with crystal violet and counted under a light microscope. Three independent biological replicates were performed.

### Collagen contraction assay

After 48-hour of knockdown with siSUV39H1, or siControl, hCASMCs were resuspended in a 10% FBS in M199 media at a density 2×10^6^ cells/mL. The SMC suspension was incorporated in a collagen solution [bovine Type I collagen solution (Cell Biolabs, CBA, #201) and as described in the protocol. Cell and the cell-collagen solution was deposed on non-treated tissue culture 24-well plates. Plates were incubated at 37°C for 1 hour to allow polymerization. After collagen polymerization, 10 % FBS culture medium is added a top each collagen gel lattice and the plates are further incubated for 48 hours. The collagen gels are then gently released from the side of the wells with a sterile syringe needle. Gel images were acquired, and maximum contraction was recorded at 6 hours. Collagen gel size change (Area of contraction) was calculated using photographs taken.

### Chromatin Immunoprecipitation (ChIP)-qPCR

ChIP was performed as previously described ^6, 13^ with minor modifications using SimpleChIP Enzymatic Chromatin IP Kit (Cell Signaling). Briefly, 5 x 10^6^ cells were cross-linked with Formaldehyde 1%, methanol free, Ultra-pure EM Grade (Polysciences Inc). DNA was sheared to 500-700 bp by sonication. After immunoprecipitation with antibodies H3K9me3 (Abcam, ab8898), H3K27Ac (Abcam, ab4729), KDM4A (Abcam, ab191433), or IgG (negative control), ChIP grade magnetic beads (CST, 9006) were used to pulldown the antibody-antigen complexes. DNA was amplified by qPCR with primers to *MYH11, MYH11* (GC repressor region), *ACTA2, CNN1, MYH10* and *LMOD1.* For Primer sequences (See Supplementary Table S3). All samples were performed in at least triplicate, from at least 3 independent experiments, and data were normalized to percentage of input.

### RNA-sequencing

Total RNA quality was determined by estimating the A260/A280 and A260/A230 ratios by nanodrop. RNA integrity was determined by running an Agilent Bioanalyzer gel, which measures the ratio of the ribosomal peaks. mRNA is purified from approximately 500 ng of total RNA with oligo-dT beads and sheared by incubation at 94°C. Following first-strand synthesis with random primers, second strand synthesis is performed with dUTP for generating strand-specific sequencing libraries. The cDNA library is then end-repaired, and A-tailed, adapters are ligated, and second-strand digestion is performed by Uracil-DNA-Glycosylase. Indexed libraries that meet appropriate cut-offs for both are quantified by qRT-PCR using a commercially available kit (KAPA Biosystems) and insert size distribution determined with the Lab Chip GX or Agilent Bioanalyzer. Samples with a yield of ≥0.5 ng/ul are used for sequencing. Samples are sequenced using 75 bp single end sequencing on an Illumina Hi-Seq 2500 according to Illumina protocols. Signal intensities are converted to individual base calls during a run using the system’s Real Time Analysis (RTA) software. Primary analysis - sample de-multiplexing and alignment to the human genome - is performed using Illumina’s CASAVA 1.8.2 software suite. Rna seq analysis was done using Partek flow and pathway analysis was done using Ingenuity pathway analysis tools.

## Results

### Methylation (SUV39H1) and demethylation expression changes over time in response to vascular injury

Reversible DNA methylation has been implicated as a central mechanism in VSMC phenotypic switching, but histone methylation, especially the role of repressive marks, has not been thoroughly investigated. The H3K9me3 mark is associated with heterochromatin, and prior studies in vitro and in vivo suggest a role for SUV39H1, the enzyme that makes this mark, in VSMC phenotype. However, the distribution of this mark and the enzymes that regulate it have not been investigated in VSMC in vivo. We assessed expression of H3K9me3, SUV39H1 and KDM4A over a time course following carotid artery ligation in wild type C57BL/6 mice. Immunostaining indicated that the H3K9me3 mark was present at low levels in a small number of cells in the uninjured vessel media but the number of positive cells and especially the signal intensity increased dramatically in the media over time post-injury (Figure 1A and 1D, supplementary figure 1A) and was strongly expressed in the neointima after 28 days (Figure 1E). Expression of SUV39H1, the enzyme that generates H3K9me3 marks, followed a similar pattern, with intense expression in the media at days 3-14 post-injury, and robust expression in the neointima after 28 days (Figure 1B, 1F and 1G and supplementary figure 1B). Conversely, the KDM4A histone demethylase was abundantly expressed in the normal media but decreased over time-post injury beginning at day 3, with lower levels in the media or neointima at 28 days (Figure 1C, 1H, 1I and supplementary figure 2A). Supplementary Figures 1 and 2 included merged images with ACTA2 staining. These data reveal a significant increase in H3K9me3 and SUV39H1, with concomitant reduction in the KDM4A enzyme that erases these marks, over time following arterial injury, with abundant H3K9 trimethylation in the neointima.

**Figure 1:**
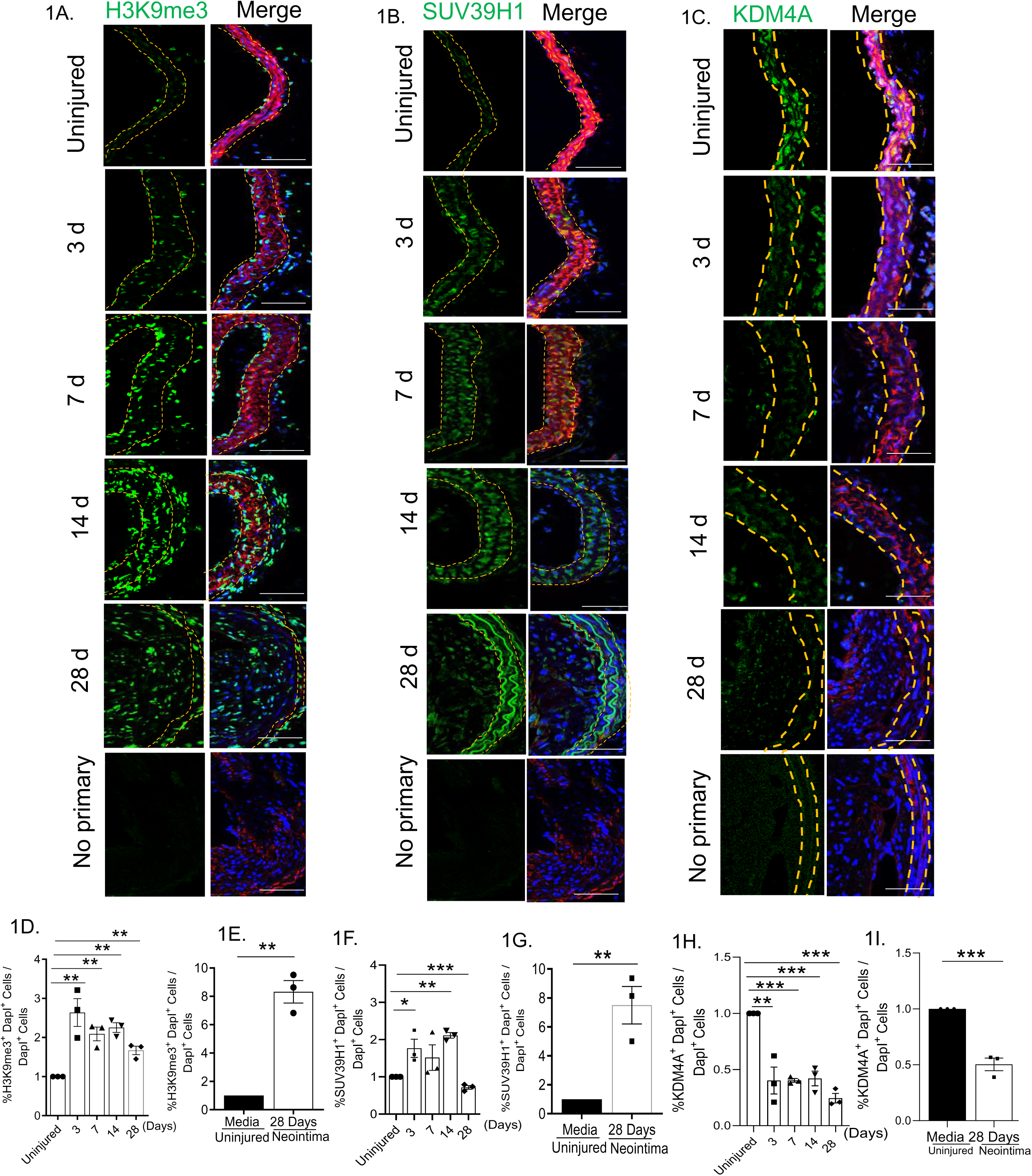
Histone H3 lysine 9 (H3K9) methylation and SUV39H1 expression changes over time in response to vascular injury. Wild-type mice were subjected to carotid artery ligation, and injured vessels were harvested at time points ranging from 3 to 28 days and compared to uninjured vessels (0 day). Cryosections were immunostained for A) H3K9me3 (green, Top left), B) SUV39H1 (green, Top Right). C) KDM4A (green, Top Right) Nuclei in all sections were additionally stained with DAPI (blue, third panel from left). A-C) Injured sections were stained with secondary antibody, smooth muscle α-actin (ACTA2), and DAPI only as a (“no primary”) negative control (bottom panels). D) Quantification of medial H3K9me3+/DAPI positive medial cells divided by total number of DAPI positive medial cells at each time point (uninjured=time 0, or days 3, 7, 14, 28 post-ligation from 2 or 3 sections per mouse from n=3 mice per time point. One-way ANOVA with Bonferroni’s post hoc test was performed. Data are expressed as mean±SEM *P<0.05, **P<0.01, ***P<0.001. E) neointimal H3K9me3, Student *t* test was performed. Data are expressed as mean±SEM *P<0.05, **P<0.01. F) Quantification of medial SUV39H1+/ DAPI positive medial cells divided by total number of DAPI positive medial cells at each time point (uninjured=time 0, or days 7, 14, 28 post-ligation from 2 or 3 sections per mouse from n=3 mice per time point. One-way ANOVA with Bonferroni’s post hoc test was performed. Data are expressed as mean±SEM *P<0.05, **P<0.01, ***P<0.001 G) Neointimal SUV39H1 positive cells as compared to uninjured (n=3) mouse carotid arteries. Student *t* test was performed. Data are expressed as mean±SEM *P<0.05, **P<0.01. H) Quantification of medial KDM4A+/DAPI positive medial cells divided by total number of DAPI positive medial cells at each time point (uninjured=time 0, or days 3, 7, 14, 28 post-ligation from 2 or 3 sections per mouse from n=3 mice per time point. One-way ANOVA with Bonferroni’s post hoc test was performed. Data are expressed as mean±SEM *P<0.05, **P<0.01, ***P<0.001. I) neointimal KDM4A, Student *t* test was performed. Data are expressed as mean±SEM *P<0.05, **P<0.01.

### Loss of function of SUV39H1 increases contractile gene expression, contractility, migration, and proliferation in smooth muscle cell (SMCs)

Given the changes in SUV39H1 and H3K9me3 during vascular remodeling, we assessed a causal role for SUV39H1 in SMC phenotypic switching using siRNA knockdown in cultured hCASMCs. SUV39H1 knockdown induced contractile gene markers of differentiation *CNN1* and *MYH11* at the mRNA level (Figure 2A) and modestly but reproducibly reduced expression of mRNAs associated with the synthetic phenotype *FN1*, *MYH10* and *OPN* (Figure 2B). The opposing effects of SUV39H1 knockdown on SMC phenotype were also noted at the protein level (Figure 2C-D). In addition to expected changes in contractile proteins, SUV39H1 knockdown led to a 50% reduction in expression of the master dedifferentiation transcription factor KLF4 (Figure 2C-D). Consistent with the observed effect on contractile protein expression, SUV39H1 knockdown increased hCASMC functional contractility in a collagen gel contraction assay (Figure 2E-F). SUV39H1 influenced other aspects of SMC phenotypic switching, with SUV39H1 knockdown significantly attenuating PDGF-BB-induced migration in Boyden Chamber migration assays (Figure 2G-H). SUV39H1 knockdown also inhibited hCASMC proliferation in a BrdU cell proliferation assay (Figure 2I-J).

**Figure 2:**
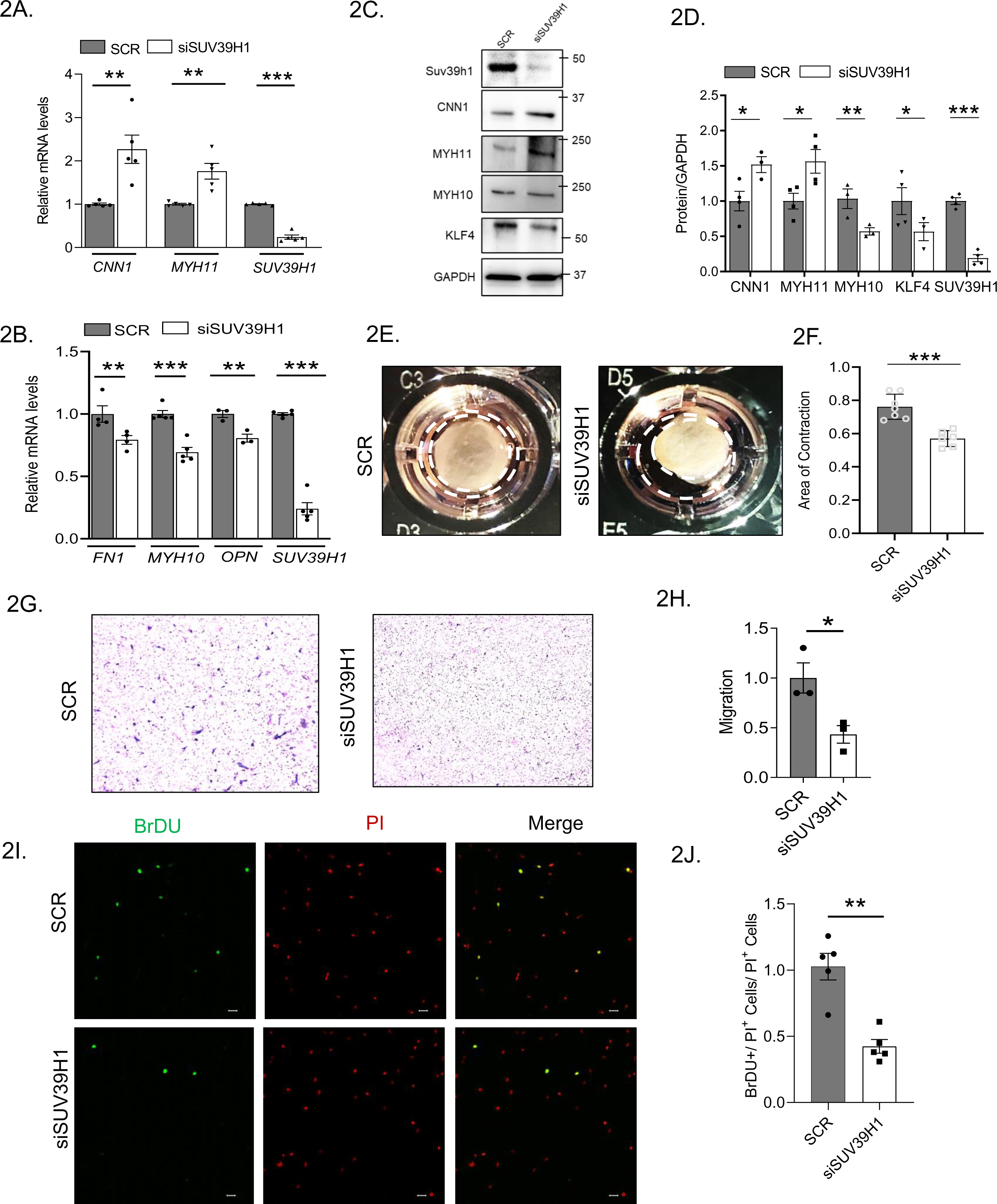
Loss of function of SUV39H1 increases contractile gene expression, contractility, migration, and proliferation in smooth muscle cell (SMCs) qPCR analysis of A) contractile genes in hCASMCs 96 hours after SUV39H1 siRNA knockdown or scrambled siRNA control and B) synthetic genes in hCASMCs 96 hours after SUV39H1 siRNA knockdown or scrambled siRNA control n=4-5 independent experiments. Student’s *t* test was performed. C, D) Western Blot analysis of key contractile, synthetic and KLF4 expression in hCASMCs following SUV39H1 knockdown. Quantification from n=4-5 independent experiments. Student’s *t* test was performed. Contractile function of hCASMCs. E) A collagen gel contraction assay revealed contractile function of hCASMCs after 96 hours of incubation. A representative collagen gel culture of both siControl and siSUV39H1 knockdown in hCASMCs is shown. F) Quantification of the area of collagen gel contraction in hCASMCs after SUV39H1 knockdown showed increased contraction compared to the scrambled siRNA control n=4-5 independent experiments. Student’s *t* test was performed. G) Boyden chamber assay to determine hCASMC migration after knockdown with siSUV39H1, or scrambled siRNA control. 48 hours after knockdown hCASMCs were seeded in the upper chamber of trans wells. PDGFBB (10ng/ml) was added to the lower Trans well chambers for 8 hours. The migrated cells were stained with crystal violet and photographed. H) 3 or 4 random fields per well were photographed and the migrated cells were counted. Quantification was normalized to average migration in the scramble control. n=3 independent experiments Student’s *t* test. I, J) Proliferation of hCASMCs after SUV39H1 siRNA knockdown or scrambled siRNA control using BrDU assay for 24 hours showed decrease in proliferation as determined by the rate of 5-Bromo-2=-deoxyuridine (BrDU) incorporation in cells. For this assay initially cells were incubated in full serum condition (M199 medium supplemented with 10% FBS) for 24 h, followed by serum starvation for another 24 h to make a synchronization of cell growth and then BrdU protocol was followed. Data are expressed as mean±SEM *\*P<0.05, **P<0.01, **P<0.001*.

### SUV39H1regulates KLF4 and PDGF-BB-induced dedifferentiation

We next assessed regulation of SUV39H1 with stimuli that induce VSMC differentiation (rapamycin) or dedifferentiation (PDGF-BB). Treatment with 10 ng/ml PDGF-BB for 24 or 48 hours decreased expression of contractile gene mRNA and proteins as expected, and increased expression of SUV39H1 (Figure 3A, 3C). Conversely, treatment with 50 nM rapamycin induced differentiation, as we have previously shown ^14, 15^, increasing contractile mRNA and protein levels (Figure 3B, 3D). Rapamycin decreased SUV39H1 protein expression (Figure 3D), but there was no change at the mRNA level (Figure 3B). By treating hCASMCs with cycloheximide over a 12-hour time course, we determined that rapamycin treatment significantly reduced the half-life of SUV39H1 protein by nearly 3 hours (Figure 3E-F).

**Figure 3:**
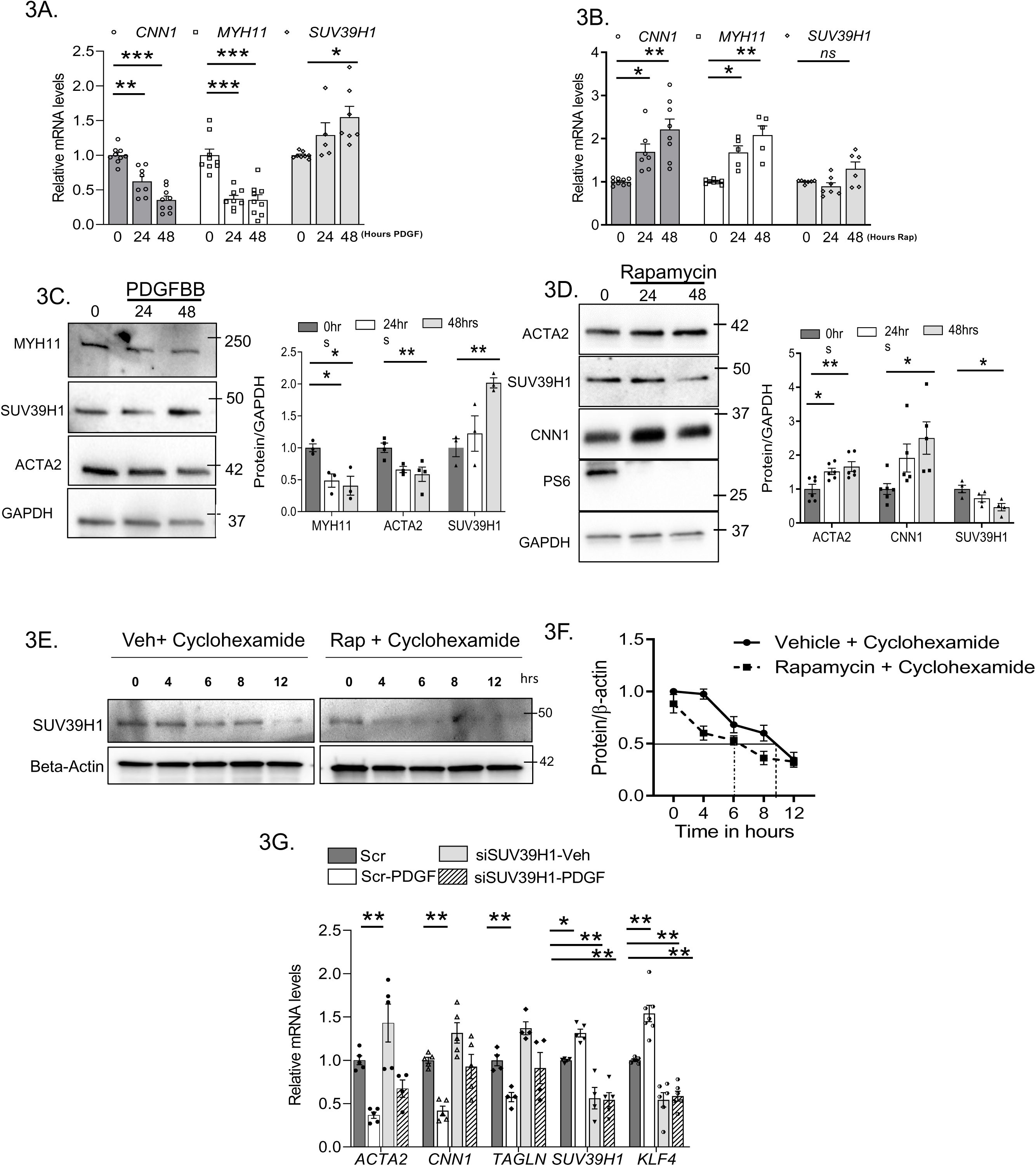
Rapamycin and PDGF differentially regulate SUV39H1expression. qPCR analysis of contractile genes and SUV39H1 in hCASMCs treated with A) 10 ng/ml PDGF-BB or B) 50 nM rapamycin treatment or vehicle for indicated time points n=5-6 independent experiments. One-way ANOVA with Bonferroni’s post hoc test was performed. Data are expressed as mean±SEM *P<0.05, **P<0.01, ***P<0.001. C) Western blot analysis of contractile proteins and SUV39H1 in hCASMCs treated with 10 ng/ml PDGF-BB or vehicle for indicated time points. D) Western blot analysis of contractile proteins and SUV39H1in hCASMCs treated with 50 nM rapamycin or vehicle for indicated time points. Quantitation from n=4-6 independent experiments. One-way ANOVA with Bonferroni’s post hoc test was performed. Data are expressed as mean±SEM *P<0.05, **P<0.01, ***P<0.001. E) hCASMCs were starved in 2.5% FBS and treated with 100 μg/mL cycloheximide (CHX). Cell lysates were harvested at various time points as indicated and subjected to western blot analysis for SUV39H1, using β-actin as loading control from n=4 independent experiments. Protein levels were quantitated, and SUV39H1 level normalized to β-actin were plotted to determine SUV39H1 protein half-life. F) Nonlinear regression analysis using 1-phase decay was performed to determine SUV39H1 protein half-life. G) qPCR analysis of contractile genes and SUV39H1 in hCASMCs after 48-hour SUV39H1 siRNA knockdown, then treated with 10 ng/ml PDGF-BB treatment or vehicle for an additional 48 hours. n=4-6 independent experiments. Two-way ANOVA followed by Sidak multiple comparisons testing was performed. Data are expressed as mean±SEM *P<0.05, **P<0.01, ***P<0.001.

To determine the role for SUV39H1 in PDGF-BB-induced dedifferentiation, we knocked down SUV39H1 in cultured hCASMCs and then treated with PDGF-BB 10 ng/ml for 48 hours. In this experiment, we achieved a 50% reduction in SUV39H1 mRNA levels which was sufficient to induce contractile gene mRNAs and induce *KLF4* expression in the absence of PDGF-BB treatment. Notably, this partial SUV39H1 knockdown completely prevented PDGF-BB induction of *KLF4,* prevented the repression of *CNN1* and *TAGLN*, and partially attenuated the *ACTA2* repression (Figure 3G). These data reveal that SUV39H1 expression is altered by stimuli that promote SMC phenotypic switching, and that SUV39H1 regulates KLF4 and mediates PDGF-BB-induced SMC dedifferentiation.

### Effect of SUV39H1 knockdown on epigenetic regulators and KLF4 expression in SMCs

Given the opposing effects of SUV39H1 on contractile and synthetic gene expression (Figures 2-3), we investigated whether SUV39H1 knockdown in cultured hCASMCs influences mRNA levels of known master regulators of SMC phenotype. SUV39H1 knockdown significantly decreased expression of KLF4 at the mRNA level, but did not significantly alter mRNA expression of SRF, myocardin, TET2, MYOCD, DNMTs, or of SUV39H2 (Figure 4A). Since SUV39H1 knockdown significantly reduced expression of KLF4 at the mRNA and protein level (Figure 2 C-D, 4A), we next determined the KLF4 mRNA half-life in hCASMCs treated over a time course with Actinomycin D and found that knock down of SUV39H1 decreased KLF4 mRNA half-life from 2 hours to 45 minutes (Figure 4B).

**Figure 4:**
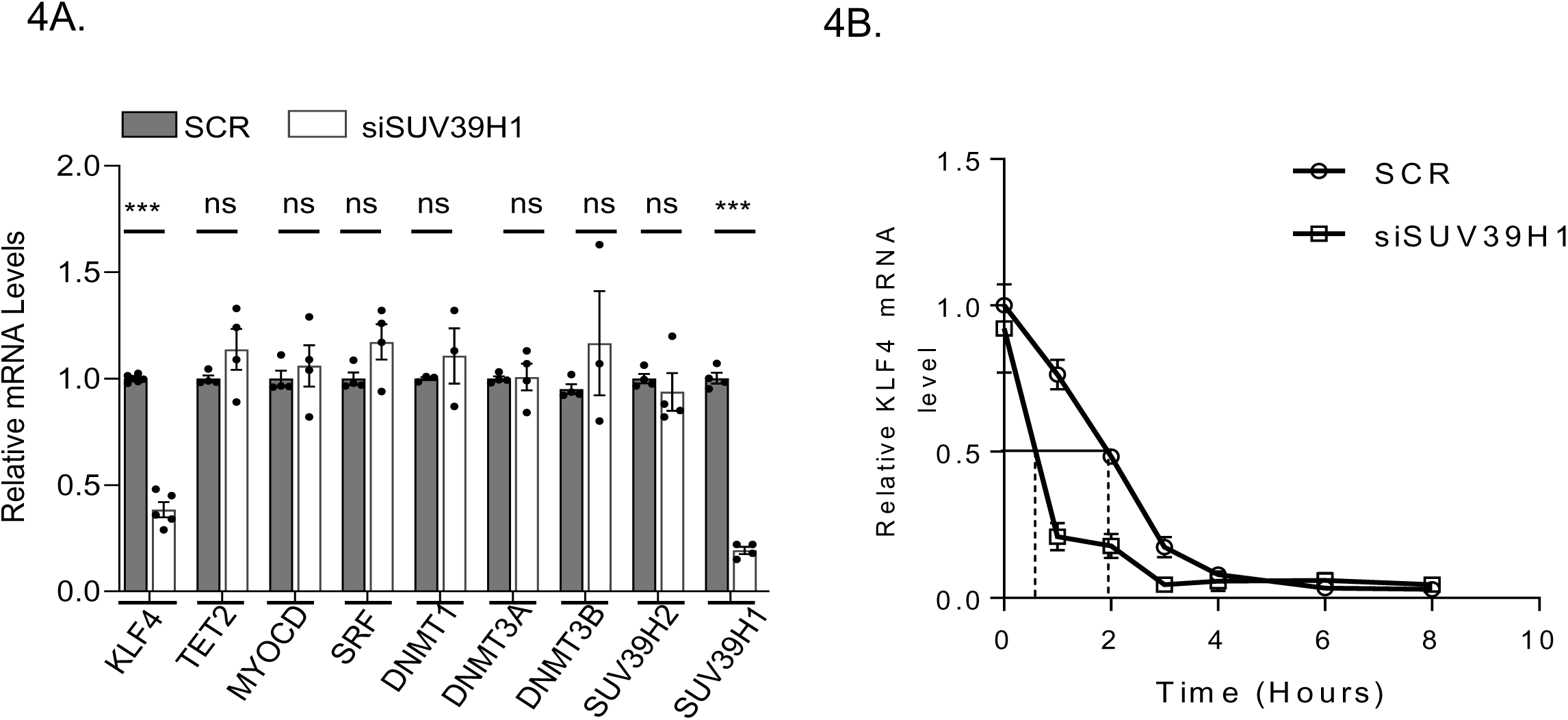
SUV39H1 affects KLF4 expression: master synthetic regulator of SMC phenotype. qPCR analysis of A) key epigenetic regulators of SMC plasticity in hCASMCs 96 hours after SUV39H1 siRNA knockdown or scrambled siRNA control. n=4-5 independent experiments. Student’s *t* test was performed. B) KLF4 mRNA half-life at time points after Actinomycin treatment in hCASMCs. SUV39H1 siRNA knockdown or scrambled siRNA control was performed. 48 hours after the knockdown SMCs were treated with actinomycin D (5 μg/mL). Quantified RT-PCR analysis of KLF4 mRNA level from a minimum of 3 independent experiments. Data are expressed as mean±SEM *P<0.05, **P<0.01, ***P<0.001.

### SUV39H1 influences both repressive H3K9me3 and activating H3K27Ac marks at contractile gene promoters

The effects of altered SUV39H1 on SMC gene expression and phenotypic switching suggested that the H3K9me3 marks that are associated with silenced heterochromatin may be malleable in VSMCs. We assessed the effects of SUV39H1 on locus-specific histone methylation at contractile gene promoters (*ACTA2, MYH11, MYH11-GC* repressor, *LMOD1*), focusing on the CArG-rich proximal promoter regions that are known to bind SRF and promote differentiation-specific gene expression ^3^. Using chromatin immunoprecipitation (ChIP-qPCR) assays, we found that SUV39H1 knockdown in hCASMCs reduced histone H3K9me3 marks at these CArG elements, as well as at the GC repressor element in the *MYH11* promoter (Figure 5A). Notably, the reduction in the repressive mark was accompanied by a concomitant decrease in DNA methylation (5mC) at these promoters (Figure 5B), and an increase in the activating mark H3K27Ac at CArG-containing regions (Figure 5C), indicating a more active chromatin state at these genes. Importantly, we demonstrated that PDGF-BB treatment induced H3K9me3 marks at these promoters, and that SUV39H1 knockdown reduced the basal H3K9me3 levels at these sites and completely prevented the PDGF-BB induction of this mark at contractile gene CArG-rich promoter regions (Figure 5D). These data reveal that PDGF induces SUV39H1-dependent repressive chromatin marks at contractile gene promoters.

**Figure 5:**
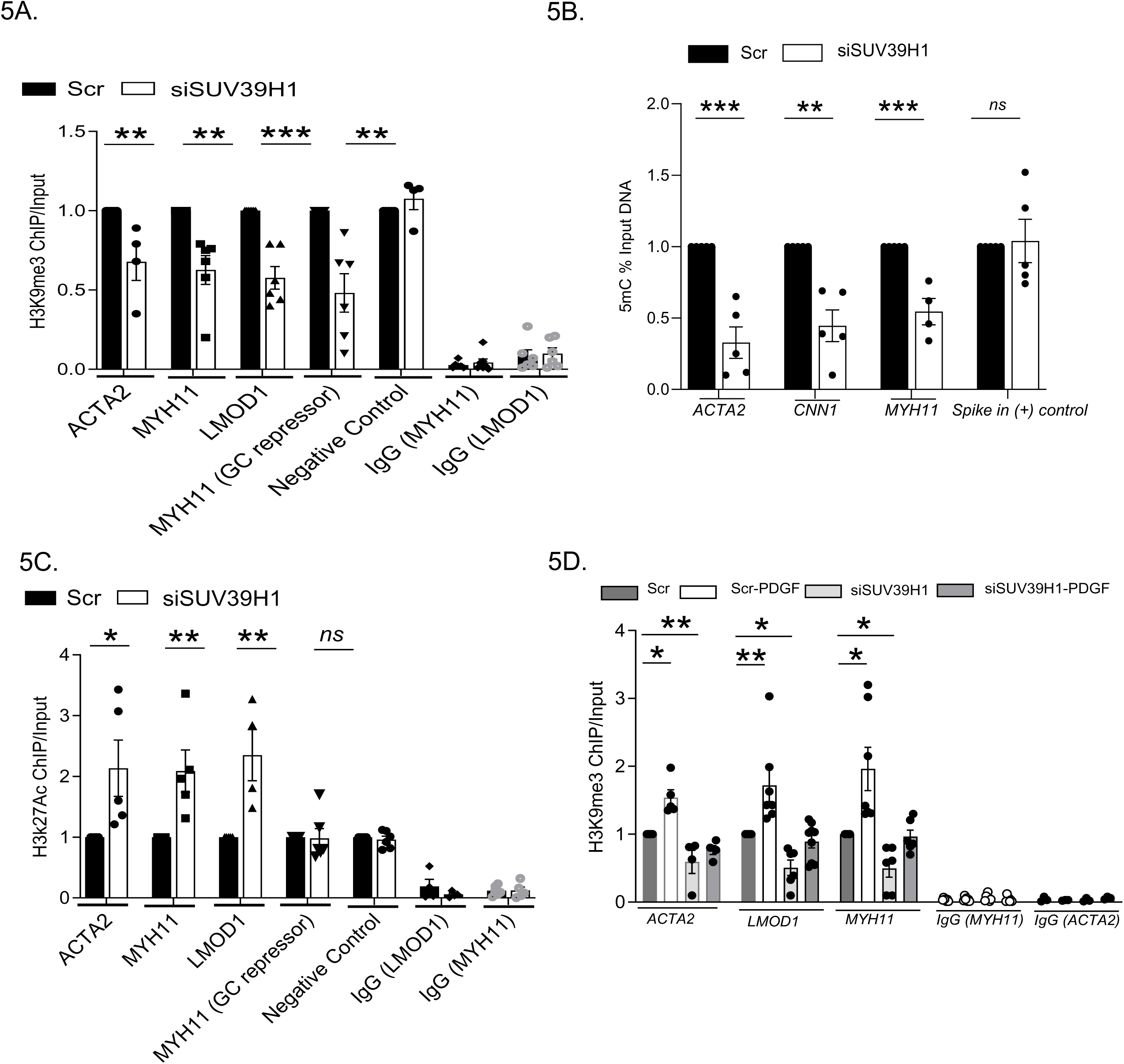
SUV39H1 regulates chromatin modification by H3K9me3 and H3K27Ac mark at contractile gene promoters. ChIP qPCR assays in hCASMCs 96 hours after SUV39H1 siRNA knockdown or scrambled siRNA control. A) ChIP assay with antibody to H3K9me3 mark at CArG-containing regulatory regions of contractile gene promoters of *ACTA2, MYH11, LMOD1,* and *MYH11 (GC repressor)* after SUV39H1 knockdown B) 5mC ChIP at contractile gene promoters of *ACTA2, CNN1* and *MYH11* from n= 4-5 independent experiments. Student *t* test: \*\**P*<0.01, \*\*\**P*<0.001. C) ChIP assay with antibody to H3K27Ac mark at CArG-containing regulatory regions of contractile gene promoters of *ACTA2, MYH11, LMOD1,* and *MYH11 (GC repressor)* after SUV39H1 knockdown. A and C, Negative control IgG immunoprecipitations were amplified with indicated primer sets. Data are presented as mean relative enrichment over input ±SEM from n=5 or 6 independent experiments. Student *t* test: \**P*<0.05, \*\**P*<0.01, \*\*\**P*<0.001. D) ChIP-qPCR assay for H3K9me3 in hCASMCs after 48-hour SUV39H1 siRNA knockdown, then treated with 10 ng/ml PDGF-BB treatment or vehicle for an additional 48 hours. n=6-7 independent experiments. Two-way ANOVA followed by Sidak multiple comparisons testing was performed. ChIP indicates chromatin immunoprecipitation.

### SUV39H1 regulation of histone demethylases

Changes in H3K9me3 marks after PDGF-BB stimulation or depletion of histone methyltransferase SUV39H1 suggests a surprising dynamic regulation of these heterochromatin-associated marks at contractile gene promoters (Figure 5 A-D). We investigated whether histone demethylases might contribute to changes in H3K9me3 at contractile genes. We assessed expression of KDM4 family histone demethylases (KDM4A, KDM4B, KDM4C and KDM4D) after SUV39H1 knockdown in hCASMCs and found that SUV39H1 knockdown induced KDM4A mRNA and protein but did not affect KDM4B or KDM4C expression (Figure 6A and 6B). KDM4D was not detected in hCASMCs. Using a KDM4A ChIP-qPCR assays, we determined that KDM4A binding to contractile gene promoters (Figure 6C) was enhanced with SUV39H1 knockdown. These data demonstrate that SUV39H1 inhibits KDM4A expression and occupancy at CArG-containing contractile gene promoters, revealing a dynamic and opposing relationship between the writer and eraser of H3K9me3 heterochromatin marks at these key regulatory regions in VSMC phenotypic switching.

**Figure 6:**
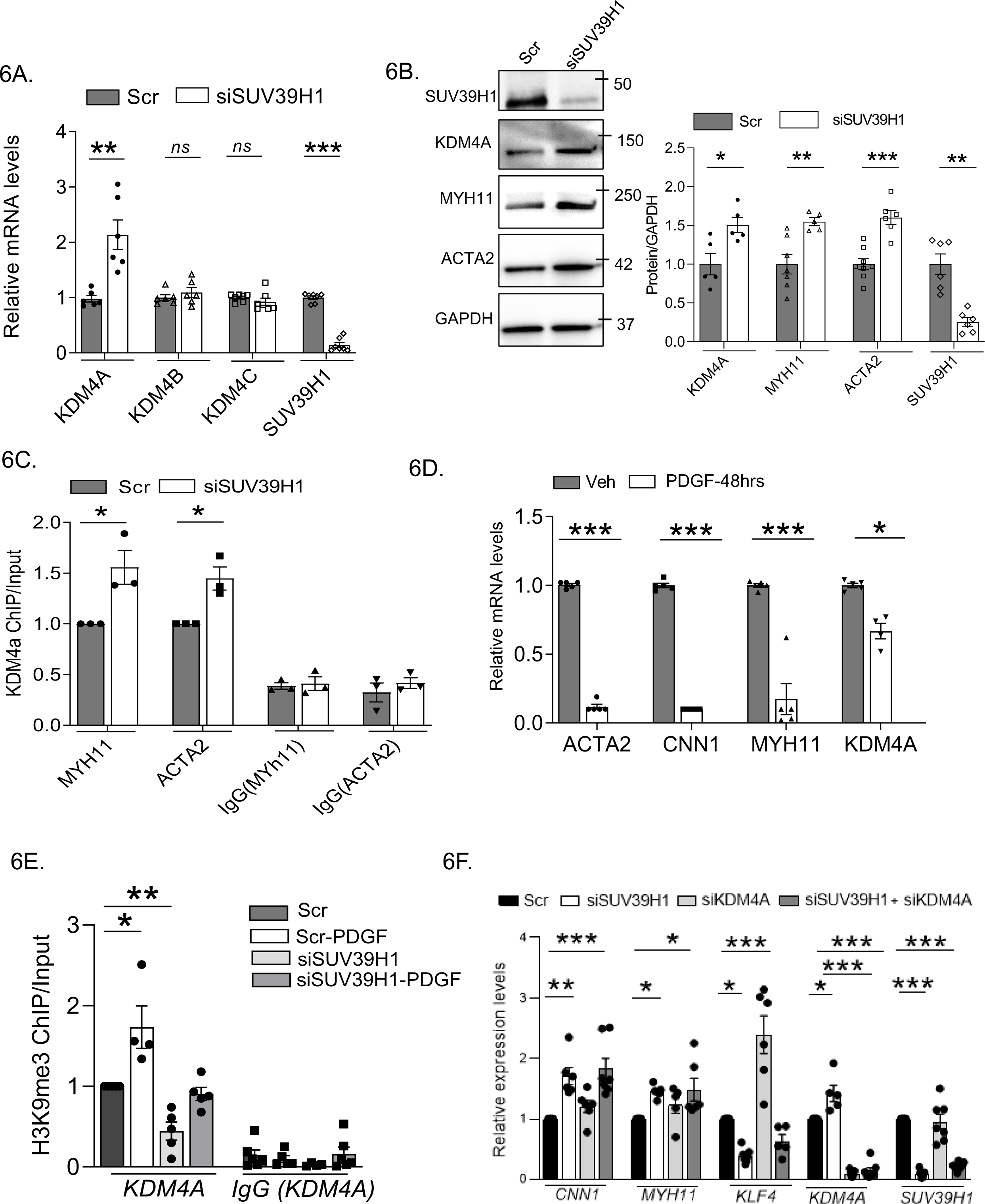
SUV39H1 regulation of histone demethylases. qPCR analysis of A) histone demethylase genes in hCASMCs after SUV39H1 siRNA knockdown or scrambled siRNA control. n=5-6 independent experiments. Student’s *t* test was performed. Data are expressed as mean±SEM *P<0.05, **P<0.01, ***P<0.001.B) Western Blot analysis of histone demethylase KDM4A and key contractile protein expression in hCASMCs following SUV39H1 knockdown. Quantification from n=5-6 independent experiments. Student’s *t* test was performed Data are expressed as mean±SEM *P<0.05, **P<0.01, ***P<0.001. C) ChIP qPCR assays for KDM4A in hCASMCs after SUV39H1 siRNA knockdown or scrambled siRNA control. Negative control IgG immunoprecipitations were amplified with indicated primer sets. Data are presented as mean relative enrichment over input. Data are represented as mean ±SEM from n=5 or 6 independent experiments. Student *t* test: \**P*<0.05, \*\**P*<0.01, \*\*\**P*<0.001. D) qPCR analysis of contractile genes and KDM4A in hCASMCs treated with 10 ng/ml PDGF-BB treatment or vehicle for indicated time points from n=4-5 independent experiments. One-way ANOVA with Bonferroni’s post hoc test was performed. Data are expressed as mean±SEM *P<0.05, **P<0.01, ***P<0.001 E) ChIP-qPCR assay for H3K9me3 in hCASMCs amplifying KDM4A proximal promoter region after 48-hour SUV39H1 siRNA knockdown, then treated with 10 ng/ml PDGF-BB treatment or vehicle for an additional 48 hours. n=5-6 independent experiments. Data are expressed as mean±SEM *P<0.05, **P<0.01, ***P<0.001. Two-way ANOVA followed by Sidak multiple comparisons testing was performed. F) qPCR analysis of contractile and KLF4 after dual knockdown of SUV39H1 and KDM4A in hCASMCs n=5-6 independent experiments. Data are expressed as mean±SEM *P<0.05, **P<0.01, ***P<0.001.Two-way ANOVA followed by Sidak multiple comparisons testing was performed.

We next assessed PDGF-BB regulation of KDM4A. We found that 48 hour PDGF treatment decreased KDM4A mRNA expression (Figure 6D). ChIP-qPCR with an anti-H3K9me3 antibody revealed that PDGF-BB treatment induced the repressive H3K9me3 at the KDM4A proximal promoter (Figure 6E). SUV39H1 knockdown reduced basal H3K9me3 at this promoter and prevented PDGF-BB from inducing H3K9 trimethylation above baseline (Figure 6E). Interestingly, we found that KDM4A knockdown reduced KLF4 expression, while SUV39H1 knockdown increased KLF4 mRNA. When both KDM4A and SUV39H1 were knocked down together, KDM4A effect predominated, with KLF4 levels remaining low. With simultaneous knockdown of both SUV39H1 and KDM4A, contractile proteins were elevated to the level seen with SUV39H1 knockdown alone (Figure 6F), consistent with a lack of the enzyme generating the repressive H3K9me3 mark and reduced KLF4 expression.

### Effect of SUV39H1 knockdown on differential gene expression

Bulk RNA-seq studies using SUV39H1 knockdown showed distinct cluster separation between control and SUV39H1 knockdown with 387 genes upregulated and 886 genes reduced (Figure 7A and B). Our RNA-seq studies also indicated significant decrease in KLF4 as well as increase in contractile gene expression (Figure 7C) with pathways as well as GO molecular functions affected shown in (Figure 7D and E).

**Figure 7.**
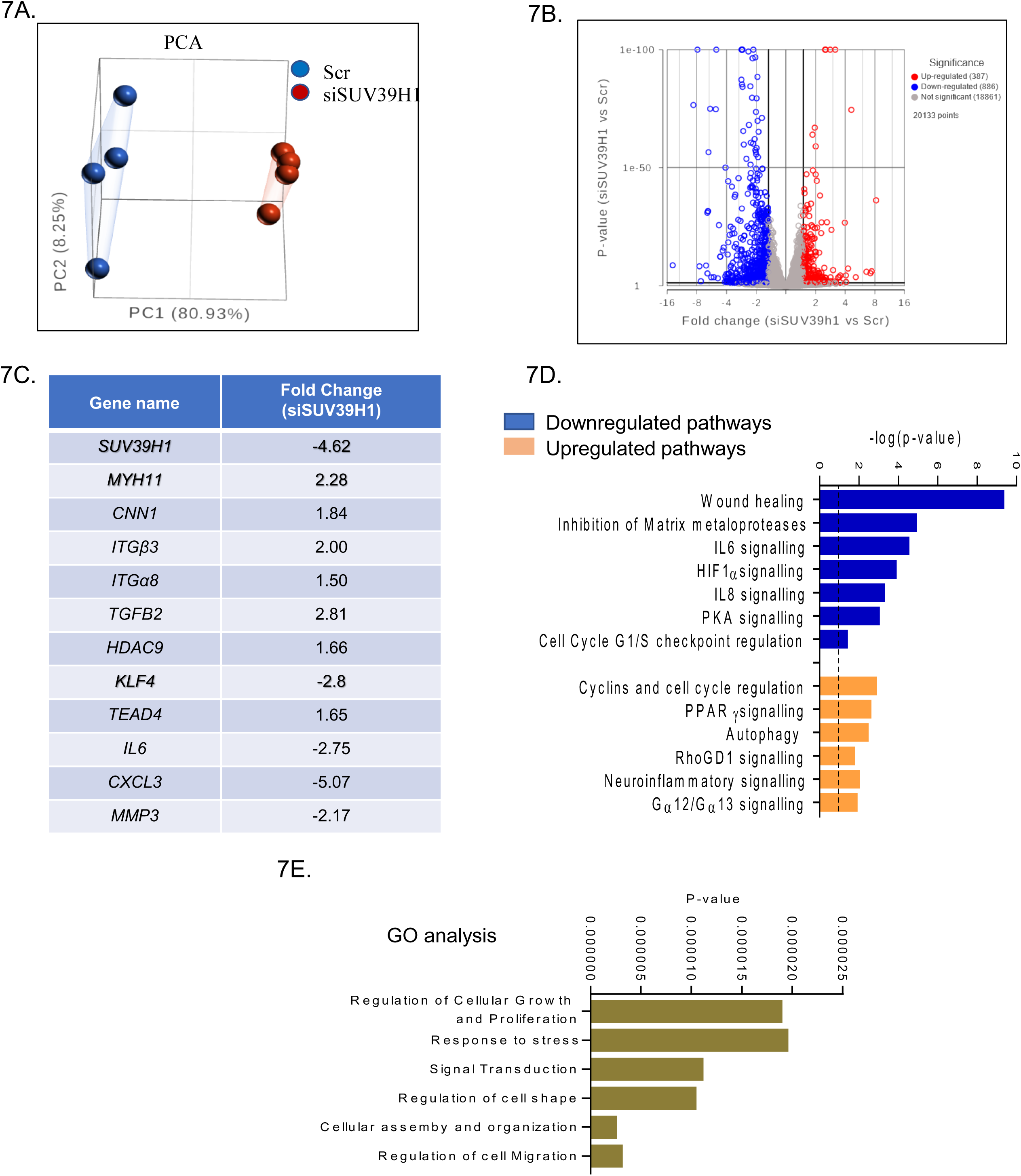
Transcriptome analyses in hCASMCs after SUV39H1 knockdown. siRNA mediated knockdown of SUV39H1 in human coronary artery smooth muscle cells (hCASMCs) were performed and bulk Rna seq analysis was done. A. PCA analysis show distinct clusters for siControl and siSUV39H1 KD SMCs. B) Volcano plot of differential regulated genes are shown. The red dots indicate upregulated genes (387) while the blue indicate downregulated genes (808) after SUV39HI knockdown. C) Genes that are differentially expressed between siControl and siSUV39H1 knockdown cells. D) Ingenuity pathway analysis (IPA) comparing siControl-versus siSUV39H1 knockdown cells. Blue bars indicate down regulated pathways while orange bars indicate upregulated pathways after SUV39H1 knockdown with indicated p values. E) Gene ontology (GO) analysis comparing siControl-versus siSUV39H1 knockdown SMCs.

**Figure 8:**
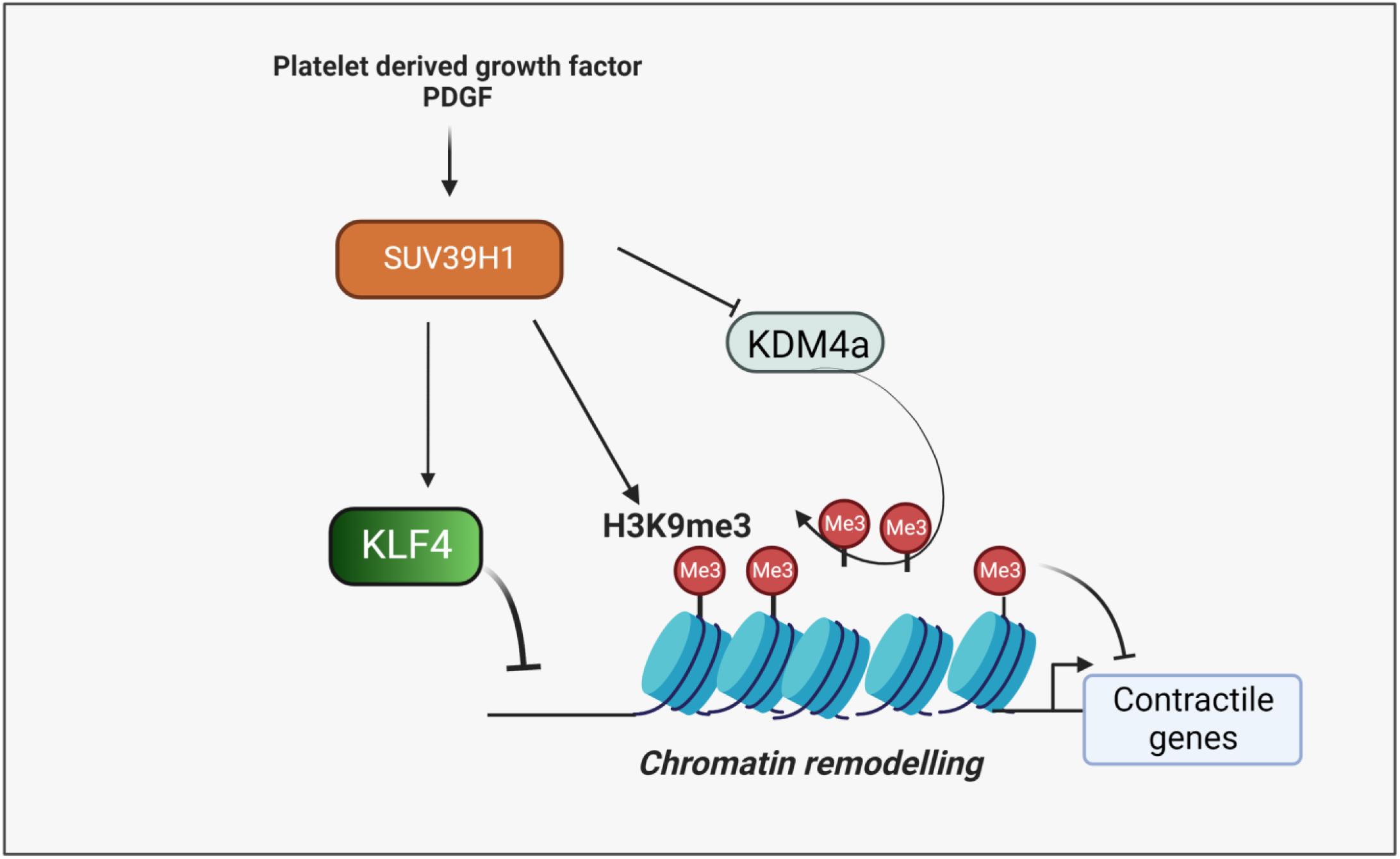
Model of the role of SUV39H1 in SMC phenotypic plasticity. Growth factor like PDGF-BB alone or when mediated through vascular injury increases histone methyl transferase SUV39H1 which in turn promotes the H3K9me3 repressive mark at contractile genes thereby promoting SMC dedifferentiation. PDGF through SUV39H1 regulates KLF4 expression, the master transcriptional regulator of dedifferentiation. Knockdown of SUV39H1 also increases eraser protein KDM4A. Thus, the net effect of PDGF is to promote H3K9me3 at contractile genes by increasing the writer (SUV) and decreasing the eraser (KDM4A).

## Discussion

We find that SUV39H1, a histone methyl transferase that generates H3K9me3, a hallmark of heterochromatin ^8^, regulates VSMC dedifferentiation. We find that SUV39H1 is induced by dedifferentiation stimuli in vitro and in vivo, and that SUV39H1-dependent reversible histone methylation plays a role in chromatin remodeling at contractile gene promoters during VSMC phenotypic switching. Our data suggest that histone methylation dynamics are regulated by SUV39H1 and KDM4A, and that regulation of H3K9me3 marks influences other chromatin modifications, including DNA methylation and activating histone acetylation (H3K27Ac).

While all differentiated cells have heterochromatin, SUV39H1 is expressed at low levels in VSMC in healthy adult mouse carotid arteries, and we detect only low level H3K9me3 staining, while expression of the histone demethylase KDM4A that removes this H3K9me3 mark is abundant in this differentiated state. This observation is notable as stem cells, where many possible gene programs are accessible, contain little heterochromatin and H3K9me3^16, 17^, suggesting an intriguing commonality between true stem cells and VSMC. Vascular injury promotes a major shift in expression of these enzymes, with a robust upregulation of the SUV39H1 writer and the H3K9me3 mark with concomitant downregulation of the eraser KDM4A. This suggests that substantial changes in chromatin repressive marks and heterochromatin reorganization accompany the phenotypic reprogramming as VSMC dedifferentiate from the contractile, quiescent phenotype to the proliferative, migratory and reparative “synthetic” phenotype in response to vascular injury. If unchecked, this repair response leads to intimal hyperplasia, which can cause restenosis in human percutaneous revascularization procedures. We note abundant expression of SUV39H1 and H3K9me3 in the mouse neointimal lesions. Prior studies employing local viral SUV39H1 knockdown, which would affect many vascular and inflammatory cell types, inhibited intimal hyperplasia ^12^. Collectively, these data suggest that this epigenetic regulator may have implications for anti-restenosis therapies.

Epigenetic modification has emerged as an important mechanism in VSMC phenotypic switching. We previously identified TET2 as a master epigenetic regulator of SMC phenotype that lies upstream of SRF, MYOCD, and KLF4, implicating reversible DNA hydroxymethylation as central to SMC plasticity ^5^. A following study further supported this relationship and noted that DNMT1 inhibits expression of TET2, documenting “yin-yang dynamics of DNA methylation” VSMC phenotypic switching ^4^ . We have also noted that distinct histone acetyltransferases regulate differentiated vs synthetic phenotype, with p300 functioning in concert with TET2 and CBP promoting SMC dedifferentiation, in part by recruiting HDACs to contractile genes ^6^. This work further noted that TET2 and p300 associate and function in concert ^6^. From these studies, a model is emerging where environmental stimuli signal to alter expression, interactions, and activities of opposing families of epigenetic regulators, leading to dynamic and substantial changes in DNA methylation and histone acetylation to promote chromatin remodeling in VSMC plasticity. This study now adds SUV39H1/KDM4A regulation of H3K9me3 to this paradigm of reversible chromatin modifications. Importantly, SUV39H1 knockdown impacted not only H3K9me3 at contractile gene promoters, but also levels of locus-specific 5mC and H3K27Ac modifications, suggesting that SUV39H1 and the H3K9me3 mark it deposits may impact the ability of other chromatin modifying enzymes to influence VSMC promoter accessibility. Further studies will help unravel whether and how specific transcription factor interactions and signaling mechanisms recruit SUV39H1 and KDM4A to VSMC promoters, and how these enzymes interact with other chromatin modifying machinery to allow for the dramatic genetic reprogramming that mediates VSMC phenotypic switching.

We have studied these epigenetic paradigms using the mTORC1 inhibitor and drug-eluting stent agent rapamycin to promote VSMC differentiation, and PDGF-BB, released at sites of vascular injury, as a dedifferentiation agent. Notably, we have found that rapamycin treatment induces expression and activity of TET2^5^ and p300 ^6^, and here we show that rapamycin inhibits SUV39H1 protein expression. PDGF-BB induces VSMC dedifferentiation, inhibiting expression of TET2^5^, p300 ^6^ and KDM4A, while increasing expression of CBP ^6^and SUV39H1. KLF4 is a master transcriptional regulator of VSMC dedifferentiation ^18^ that is induced by PDGF-BB and by SUV39H1. As the direct enzymatic activity of SUV39H1 promotes gene repression, it is likely that it enhances KLF4 mRNA half-life by inhibiting a protein or perhaps a miRNA that normally destabilizes this message. This effect on KLF4 stabilization likely works in concert with the epigenetic changes by SUV39H1 to coordinately induce VSMC dedifferentiation.

Interestingly, SUV39H1 has been implicated in the epigenetic metabolic memory of VSMC exposure to diabetes. VSMC isolated from diabetic db/db mice had persistently reduced H3K9me3 modifications at the promoters of pro-inflammatory genes, leading to increased inflammatory gene expression, even when cultured in normal media. Exposure to hyperglycemia downregulated SUV39H1 expression ^13,10^, which was mediated by a miRNA mechanism ^12,9^. KDM4A was found to be increased in SMC in response to high glucose, with reduced H3K9me3, but increased SMC proliferation and migration that were inhibited with KDM4 inhibition, and local vascular global inhibition of KDM4A attenuated intimal hyperplasia in diabetic rats ^19^. We now demonstrate that PDGF-BB and rapamycin can alter SUV39H1 expression in VSMC phenotypic switching. Signaling to H3K9me3 enzymes and other regulatory factors may differ with distinct stimuli. It will be of interest to determine in future studies mechanisms that allow for persistent versus reversible changes in SUV39H1 expression, specific gene targeting, and H3K9me3 marks.

H3K9 demethylation is mediated by members of the JmJd domain family. KDM4A (previously Jumonji domain-containing 2A (JMJD2A)), catalyzes H3K9me3 demethylation ^20^. KDM4 demethylates H3K9 and facilitates reprogramming in embryos ^21^. JMJD2A has been extensively studied in cancer development, transcriptionally regulates cancer-related genes including cell-cycle, proliferation, and inflammation^22^. Interestingly, in embryonic stem cells, H3K9me3 silences developmental regulatory genes and functions as a co-repressor of Oct3/4. In turn, Oct3/4 positively regulate the expression of two key demethylases, KDM3A and KDM4C, which remove the heterochromatin-associated H3K9me3, ensuring the maintenance of embryonic stem cell renewal. These opposing actions regulating histone H3 methylation and demethylation in stem cells are concordant with our new observations of a role for dynamic H3K9me3 regulation in VSMC plasticity. Further studies of the roles of this mark, regulatory enzymes, and heterochromatin organization in VSMC may provide exciting new insights into the mechanisms underlying VSMC phenotypic plasticity in reparative vascular remodeling and in cardiovascular disease.

## Subject Terms

Smooth muscle cells

Epigenetics and chromatin remodeling Neointima formation

histone methyl transferase,

histone demethylase JMJDs/KDM4 Phenotypic switch,

Basic science research

## Nonstandard Abbreviations and Acronyms

hCASMCs: Human coronary artery smooth muscle cells
PDGF: Platelet-derived growth factor
mTORC1: Mammalian target of rapamycin complex 1
ChIP-qPCR: Chromatin Immunoprecipitation-quantitative
PCR KDMs: histone lysine demethylases
SUV39H1: Suppressor Of Variegation 3-9 Homolog 1

---

**Supplementary Figure 1.**
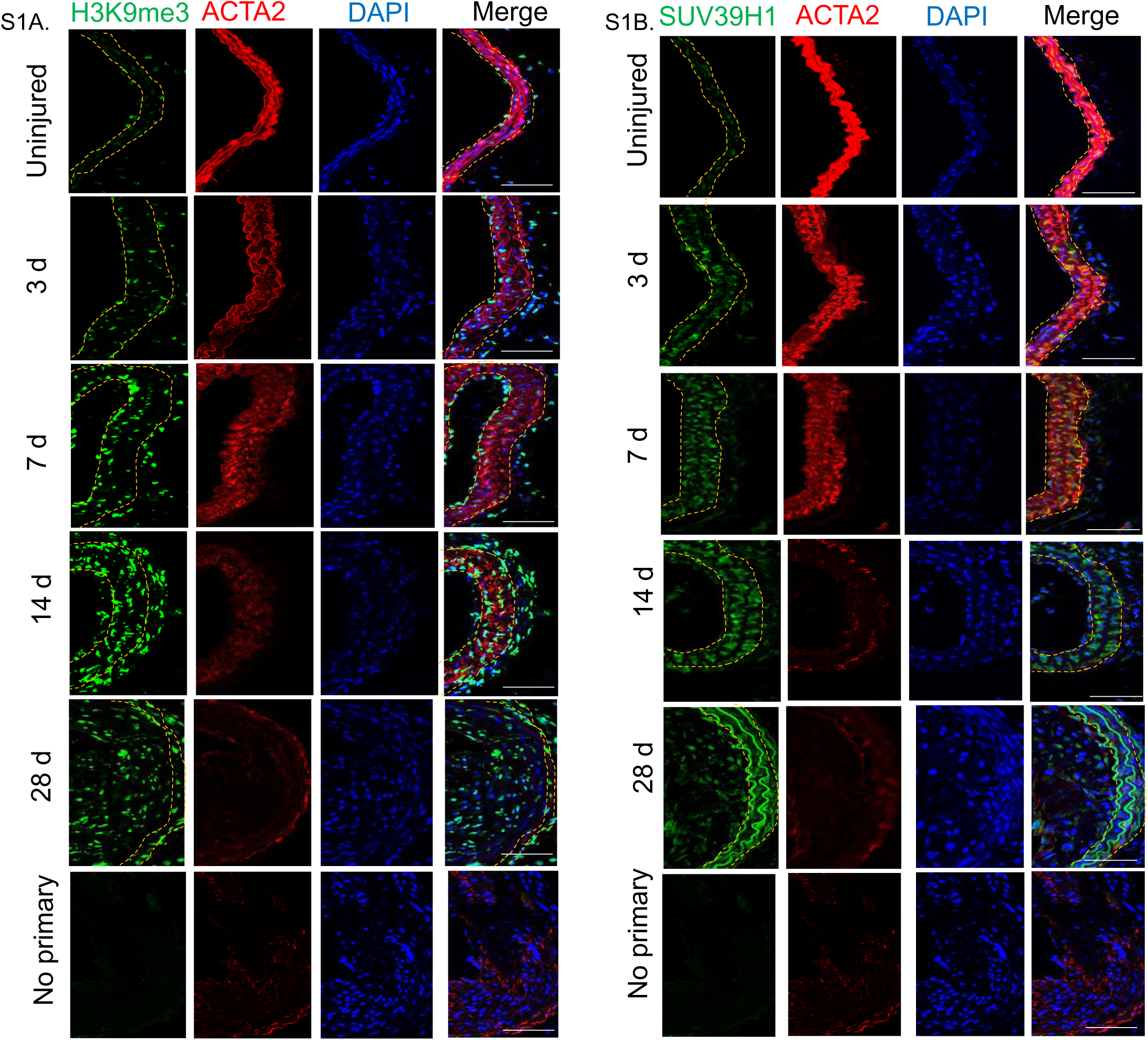

**Supplementary Figure 2.**
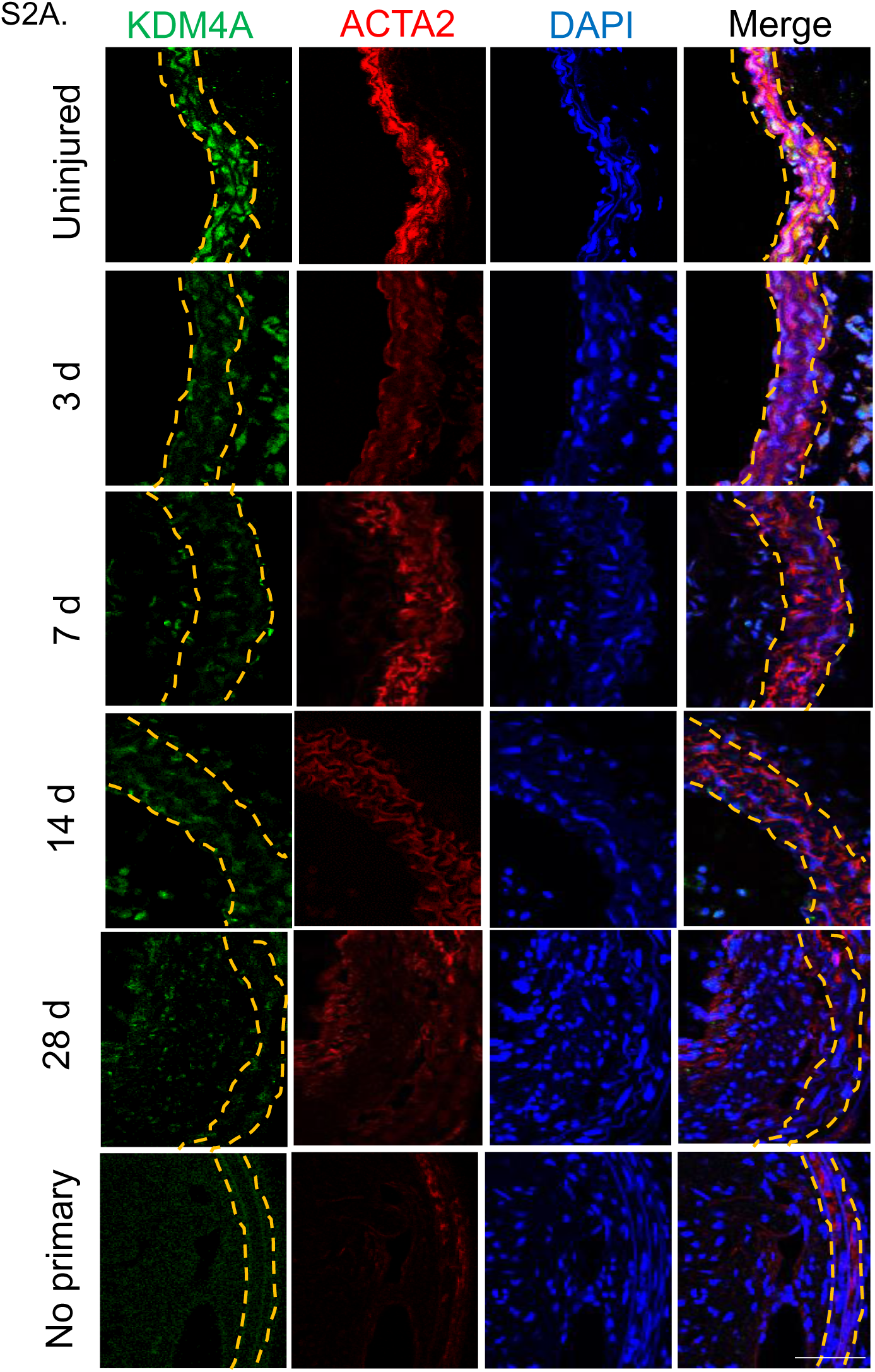

## Notes

### Competing Interest Statement

The authors have declared no competing interest.

