## Supplementary data for "SUV39H1 mediated regulation of KLF4 and KDM4A coordinate smooth muscle cell phenotypic plasticity"

**Extended methods:**

**Quantitative PCR**

Total RNA was isolated from using the Qiagen RNeasy kit, and 1 mg of RNA was reverse transcribed using All-In-One 5X RT Master Mix cDNA synthesis kit (ABM). qPCR was performed using BlasTaq™ 2X qPCR Master Mix (ABM, G891). Primer sequences are listed in Supplementary Table S1.

**Western blot**

hCASMCs were cultured as described [4, 8]. Before treatment, cells were serum starved for 24 hours in media with 2.5% FBS for Rapamycin treatment or with 0.5% FBS for PDGF treatment. Cells were then treated with Rapamycin, PDGF-BB or vehicle for additional indicated time points. Cells were lysed in RIPA buffer with protease inhibitors and protein concentration was determined using BCA assay. 10-20µg of proteins was loaded in each lane of 4-15% SDS-PAGE gels, transferred to Immobilon PVDF membranes (Millipore), blocked with 5% nonfat dry milk, washed in Tris-buffered saline, 0.1% Tween 20 (TBS-T) and probed with primary antibodies (See Supplementary Table S2) overnight at 4°C. Membranes were incubated with HRP-conjugated secondary antibodies (Invitrogen) (See Supplementary Table S2), washed in TBS-T, developed with Supersignal West Pico Plus Chemiluminescent Substrate or West Femto Maximum Sensitivity Substrate (ThermoScientific) and analyzed with the G:BOX imaging system (Genesys).

**Proliferation assay**

Proliferation was assessed as previously described with minor modifications. SMCs were treated with either siRNA control or siSUV39H1 for 24 hours and were then trypsinized and cultured overnight on culture slides. The next day, the cells were washed with PBS and serum starved overnight in M199 media supplemented with 0.5% FBS to make a synchronization of cell growth.. Cells were then cultured for 24 hours in complete medium for 24 hours. For the final 10 hours of this incubation, 10 µg/ mL BrdU (Millipore Sigma) was added to the cells. Slides were fixed in 4% paraformaldehyde for 30 minutes, rinsed in 0.3% Tris and 1.5% glycine in water for 15 minutes, incubated in 2N HCl for 30 minutes at 37°C, washed with 0.1 M boric acid, and then blocked with 0.5% NGS in PBS-T for 1 hour. SMCs were stained with rat anti-BrdU primary antibody (1:100, Bio-Rad catalog MCA2060) in 0.5% NGS in PBS-T for 1 hour, washed 3 times in 0.5% Tween 20 in PBS, and then incubated with goat anti-rat secondary antibody conjugated to Alexa Fluor 488 (1:500, Molecular Probes catalog A-11006) and PI (1:500, Millipore Sigma) in 0.5% NGS in PBS-T for 1 hour. Finally, slides were washed 3 times in 0.5% Tween 20 in PBS and mounted on slides using fluorescence mounting medium (Dako). Proliferation was calculated as the percentage of total PI+ SMCs that were BrdU+. For each condition, at least 3 fields of view were quantitated.

**Methyl (MeDIP)-qPCR:**

MeDIP was carried out using the protocol according to manufacturer's instructions. Briefly enrichment of methylated DNA was performed using the Me-DIP kit (Diagenode # Cat C02010031). A total amount of 1µg of Genomic DNA was sheared to 200 – 500 bp using the Covaris sonicator. Methylated DNA was captured by incubation with 5-mC antibody to magnetic beads overnight at 4°C. The beads were washed, and DNA eluted. The methylated DNA and input fractions were analyzed by qPCR to confirm enrichment of the methylated gene.

**Supplementary Table S1**

| Species | Gene | Forward primer | Reverse primer |
|---|---|---|---|
| Human | *18s* | TAACGAACGAGACTCTGGCAT | CGGACATCTAAGGGCATCACAG |
| Human | *TET2* | TGGCAAACATTCAGCAGCAC | TTGCCCTCAACATGGTTGGT |
| Human | *KLF4* | CATCTCAAGGCACACCTGCGAA | TCGGTCGCATTTTTGGCACTGG |
| Human | *CNN1* | ATGTCCTCTGCTCACTTC | ATACTTCTGGGCCAGCTTGTT |
| Human | *ACTA2* | CTATGCCTCTGGACGCACAACT | CAGATCCAGACGCATGATGGCA |
| Human | *MYH11* | CCTTGAGGAGAGGATTAGTGA | TTCCTTCTTTAGCCGCACTTC |
| Human | *TAGLN* | AAGCGCAGGAGCATAAGAGG | CTCTGTTGCTGCCCATCTGA |
| Human | *OPN* | AGTTTCGCAGACCTGACATCCAGT | TTCATAACTGTCCTTCCCACGGCT |
| Human | *FN1* | AAACCAATTCTTGGAGCAGG | CCATAAAGGGCAACCAAGAG |
| Human | *MYH10* | ATGAGCGTCGACACGCGGAC | TGGCACGCGTCGCTTCTTCT |
| Human | *SUV39H1* | CCGCCTACTATGGCAACATCTC | CTTGTGGCAAAGAAAGCGATGCG |
| Human | *SRF* | TCACCTACCAGGTGTCGGAGTC | GTGCTGTTTGGATGGTGGAGGT |
| Human | *LMOD1* | GAGGCCATGCTCAACTTCTG | CTCTCCATTCTTGGCATCTG |
| Human | *SUV39H2* | ATTGATAACCTCGATACTCGTCTT | TCTCCAGAACCTTTCATTTGATAA |
| Human | *MYOCD* | CTTATTGAAAGCGGAGAAATG | TGGGTATCTTTGGGACTTTTTG |
| Human | *DNMT1* | ACCATGACAGGAAGAACGGC | CTTCCACGCAGGAGCAGA |
| Human | *DNMT3A* | GCTGGGAGTCCAGCCTCCGT | CCAGCCACTCGTCCCGCTTG |
| Human | *DNMT3B* | TAACAACGGCAAAGACCGAGGG | TCCTGCCACAAGACAAACAGCC |
| Human | *KDM4A* | GAGGAAGACTGCTGCTTATGCTC | TCACATCCACTGGACTTCTTTCA |
| Human | *KDM4B* | TGCAGTCCCTGAGGTACGATT | CCCGACACTCTCTTCATACGC |
| Human | *KDM4C* | CCTTCAGCAGAGACACATTTCCT | TCCAAGATACTTTGCCCCATAGA |

**Table S1. Primer pair sequences for qRT-PCR.** This table shows primer pairs utilized for qRT-PCRanalysis in the study with human RNA samples.

**Supplementary Table S2**

| Species | Gene | Forward primer | Reverse primer |
|---|---|---|---|
| Human | *ACTA2* | CGGCCACCCAGATTAGAG | CTGCTCTCCTCCCACTTGC |
| Human | *LMOD1* | AGTACTAGCCAGGCACTTCA | GGAGAAACCGGGAAATCTCTTT |
| Human | *CNN1* | GGTGGAAGAAGGCTGGTCTC | CCTTGTGATGTCAGGCCCTT |
| Human | *MYH11* | GAGATGGCAAGTTGGGAAAA | GCTGTGGTGGATCTGACTT |
| Human | *KDM4A* | GAGAGCCTCCCTCATGGTCT | GGCCTTGGAAACCCCTCATT |
| Human | *MYH11-GC repressor* | GGGCGGGAGACAACCAAAA | GGAAGGCCACTCGGCACCAT |

**Table S2. ChIP Primer pair sequences for ChIP-qPCR.**

| *Target* | *Vendor source #Cat number* | *Dilution* | *Application* |
|---|---|---|---|
| ACTA2 | Sigma, A2547 | 1:1000 | W:B |
| CNN1 | Thermo Scientific, RM-2102-S0 | 1:1000 | W:B |
| H3K9me3 | Abcam: Ab8898 | 1:500, 1:50 | IF, ChIP |
| SUV39H1 | Invitrogen, 702443 | 1:500 | W:B, IF |
| KDM4A | Abcam, Ab191433 | 1:1000, 1:500, 1:50 | W:B, IF, CHIP |
| H3K27Ac | Abcam, ab4729 | 1:250, 1:50 | IF, ChIP |
| MYH11 | Biomedical Technologies, BT-562: Sigma, M7786 | 1:1000 | W:B/IF |
| MYH10 | Abcam, Ab230823 | 1:1000 | W:B |
| LMOD1 | Proteintech, 15117-1-AP | 1:1000 | W:B |
| GAPDH | CST, 2118S | 1:1000 | W:B |
| KLF4 | CST, 12173S, Abcam, ab215036 | 1:1000 | W:B |
| p-S6 ribosomal protein (Ser240/244) | CST, 5364 | 1:1000 | W:B |
| BrDU | Bio-Rad, MCA2060 | 1:500 | IF |
| α-mouse-HRP | Thermo Scientific, 31450 | 1:2000 | W:B |
| α-rabbit-HRP | Thermo Scientific, 31460 | 1:2000 | W:B |
| α-goat-HRP | Thermo Scientific, 31402 | 1:2000 | W:B |
| α-rabbit-alexa488 | Invitrogen, A11008 | 1:2000 | IF |

**Supplementary Table S3**

**Primary antibodies and secondary antibodies:**

**Supplementary Figures:**

**Supplementary Figure 1: Histone H3 lysine 9 (H3K9) methylation and SUV39H1 expression changes over time in response to vascular injury**

Wild-type mice were subjected to carotid artery ligation, and injured vessels were harvested at time points ranging from 3 to 28 days and compared to uninjured vessels (0 day). Cryosections were immunostained for A) H3K9me3 (green, Top left), B) SUV39H1 (green, Top Right). Nuclei in all sections were additionally stained with DAPI (blue, third panel from left) and ACTA2 (shown in red). A-B) Injured sections were stained with secondary antibody, smooth muscle α-actin (ACTA2), and DAPI only as a ("no primary") negative control (bottom panels).

**Supplementary Figure 2: Histone demethylation (KDM4A) expression changes over time in response to vascular injury**

Wild-type mice were subjected to carotid artery ligation, and injured vessels were harvested at time points ranging from 3 to 28 days and compared to uninjured vessels (0 day). A) Cryosections were immunostained for KDM4A (green). Nuclei in all sections were additionally stained with DAPI (blue, third panel from left) and ACTA2 (second panel from right red). Injured sections were stained with secondary antibody, smooth muscle α-actin (ACTA2), and DAPI only as a ("no primary") negative control (bottom panels).
